# High complexity and degree of genetic variation in *Brettanomyces bruxellensis* population

**DOI:** 10.1101/826990

**Authors:** Jean-Sébastien Gounot, Cécile Neuvéglise, Kelle C. Freel, Hugo Devillers, Jure Piškur, Anne Friedrich, Joseph Schacherer

**Author notes:** Corresponding authors (A.F.), (J.S).

## Abstract

Genome-wide characterization of genetic variants of a large population of individuals within the same species is essential to have a deeper insight into its evolutionary history as well as the genotype-phenotype relationship. Population genomic surveys have been performed in multiple yeast species, including the two model organisms, *Saccharomyces cerevisiae* and *Schizosaccharomyces pombe*. In this context, we sought to explore an uncharacterized yeast species, *Brettanomyces bruxellensis*, which is a major cause of wine spoilage but also can contribute to the specific flavor profile of some Belgium beers. We have completely sequenced the genome of 53 *B. bruxellensis* strains isolated worldwide. The annotation of the reference genome allowed us to define the gene content of this species. As previously suggested, our genomic data clearly highlighted that genetic diversity variation is related to ploidy level, which is variable in the *B. bruxellensis* species. Genomes are punctuated by multiple loss-of-heterozygosity regions while aneuploidies as well as segmental duplications are uncommon. Interestingly, triploid genomes are more prone to gene copy number variation than diploids. Finally, the pangenome of the species was reconstructed and was found to be small with few accessory genes compared to *S. cerevisiae*. The pangenome is composed of 4,923 core ORFs and 303 ORFs that are variable within the population. All these results highlight the different trajectories of species evolution and consequently the interest of establishing population genomic surveys in more populations.

## Introduction

The yeast species *Brettanomyces bruxellensis* (anamorph *Dekkera bruxellensis*) is a distant relative of *Saccharomyces cerevisiae* since lineages diverged more than 200 million years ago. Interestingly, these species share characteristics that have been independently acquired during evolution, such as the metabolic ability to produce ethanol in the presence of oxygen and excess of glucose (Schifferdecker, et al. 2014). Both species also share the capacity to efficiently catabolize the produced ethanol, their corresponding life style being described as ‘make-accumulate-consume (ethanol)’ (Thomson, et al. 2005). This strategy allows *B. bruxellensis*, which is associated with several human fermentation processes, to survive and develop in harsh and limiting environmental conditions. Until now, this yeast species has exclusively been isolated from anthropized niches.

The impact of *B. bruxellensis* on the fermentative processes is, however, diverse. As an example, it has a positive contribution to brewing of Belgium Lambic and Gueuze beers, but it is also well known as a major player leading to wine-spoilage by producing odorant molecules (volatile phenol) described as having barnyard or horse sweat characteristics (Conterno, et al. 2006). It has also been found associated with other food or industrial processes, such as kombucha or bioethanol, for which its contribution is still unclear (Beckner, et al. 2011; Teoh, et al. 2004). The opposite contributions of *B. bruxellensis* in fermentation processes raised a growing interest in this species, and high phenotypic variability regarding for example sugar metabolism or nitrogen source utilization has been highlighted among the isolates in various studies (Borneman, et al. 2014; Crauwels, et al. 2015; Crauwels, et al. 2017).

The observed phenotypic variability is undoubtedly, at least in part, related to its genomic plasticity, which is another peculiarity of this species that defines it as a good model at the evolutionary level. Indeed, the first proof of *B. bruxellensis* high genomic variability was provided through the comparison of electrophoretic karyotypes from different isolates that showed extensive chromosomal rearrangements (Hellborg and Piskur 2009), indicating rapid evolution at the intraspecific level. Whole genome sequencing of a small number of isolates revealed variation of the ploidy level among isolates with some triploid individuals derived from an allotriploidization event involving a moderately heterozygous diploid and a more distantly related haploid (Borneman, et al. 2014). The level of ploidy is supposed to be linked to the substrate of isolation and geographical distribution (Albertin, et al. 2014). Traditionnally, genomic variability has essentially been restricted to the exploration of small regions of the genomes (Conterno, et al. 2006; Crauwels, et al. 2014; Avramova, et al. 2018; Agnolucci, et al. 2009; Curtin, et al. 2007; Vigentini, et al. 2012). More recently, the complete genomes of a limited number of strains have been sequenced and assembled (Borneman, et al. 2014; Crauwels, et al. 2014; Curtin, et al. 2012; Piskur, et al. 2012; Olsen, et al. 2015). Comparison of these genomes suggested that polyploid and hybridization events might play a significant role in the evolution of this species (Borneman, et al. 2014). The most recent population survey was performed on a large collection of almost 1,500 isolates based on 12 microsatellite regions and showed that the population is structured according to ploidy level, substrate of isolation, and geographical origin of the strain (Avramova, et al. 2018).

With the currently available sequencing technologies, it is now possible to explore the intraspecific variability of a species at the genome-wide level. Such population genomic studies have been performed on multiple yeast species, including *S. cerevisiae* (Peter, et al. 2018; Skelly, et al. 2013; Bergström, et al. 2014; Strope, et al. 2015; Almeida, et al. 2015; Zhu, et al. 2016; Gonçalves, et al. 2016, Gallone, et al. 2016) and *Schizosaccharomyces pombe* (Fawcett, et al. 2014; Jeffares, et al. 2015) but also non-model yeast species (Leducq, et al. 2016; Carreté, et al. 2018; Hirakawa, et al. 2015; Ford, et al. 2015; Ropars, et al. 2018; Friedrich, et al. 2015; Ortiz-Merino, et al. 2018), granting better insights into their respective evolutionary histories as well as genotype-phenotype relationships. Here, we conducted a population genomic survey of *B. bruxellensis* based on whole genome sequencing data. In total, 53 worldwide collected isolates were completely sequenced using Illumina short-read technology. The data shows a high complexity of genetic variation among isolates. This species displays a variable level of nucleotide diversity, heterozygosity and copy number variants, depending on the ploidy of the isolates. This dataset offers a first view of the genomic variants at a genome-wide scale within *B. bruxellensis* and overall, provides insights into the evolutionary history of the species as well as the identification of genetic contents linked to subpopulation adaptations.

## Results and discussion

In order to survey genome-wide variability within the *B. bruxellensis* species, we gathered a collection of 53 isolates from diverse origins (Table S1), with a large part of our collection being wine-related. This collection is representative of all five major clusters previously defined using microsatellites based on almost 1,500 isolates (Avromava, et al. 2018). These strains were mostly isolated in Europe (*e.g.* Belgium, Italy, Spain) but also South Africa, Australia and Chile. We sequenced the genomes of all strains with a short-read sequencing strategy at a 40-fold coverage at least, with a mean of 98-fold coverage. The genetic diversity was explored through the comparison of the isolate sequences with a recently published reference sequence of the species (Fournier, et al. 2017), as well as through the construction of genome assemblies in order to define the gene repertoire of the species.

### Gene content of the *B. bruxellensis* genome

The recent release of a highly contiguous reference assembly of *B. bruxellensis* (Fournier, et al. 2017) opens the way to a genome-wide exploration of intraspecific variability. However, functional analyses of the genomic variability require genetic elements to be located on the genome sequence, which led us to carry out a complete annotation of this assembly. At first, a total of 5,226 protein-coding genes were predicted, among which 1,427 were interrupted by frameshifts or stop codons probably due to sequencing or assembly errors linked to the heterozygous diploid state of the sequenced isolate. This high proportion of out-of-frame genes led us to refine the reference sequence. This step was performed by retrieving the sequences of the concerned genes in an independent assembly constructed with Illumina reads and scaffolded with Redundans (Pryszcz, et al. 2016). This procedure allowed us to infer 872 in-frame gene sequences in the initial assembly leading to a total of 4,671 in-frame protein coding genes (90%) within the genome. The remaining 555 out-of-frame genes were considered to be pseudogenes.

Publicly available RNA-seq data (Piskur, et al. 2012) allowed for the identification of 509 introns in 472 genes, *i.e*. 9% of the genes, with 24 introns located in UTR. In total, 99 tRNA genes with 39 different anticodons were listed. The 26 transposable elements found in the UMY321 reference genome belong to the LTR retrotransposons. Most of them are closely related to *Debaryomyces hansenii* Tdh5 (Neuvéglise, et al. 2002). In addition to the intact and degenerate copies, at least 96 solo LTR were detected, sometimes grouped into large regions of up to 40 kb that may correspond to centromeres as reported for *D. hansenii* (Lynch, et al. 2010).

### Genetic diversity variation is related to ploidy level within *B. bruxellensis*

The mapping of the Illumina paired-end reads on the UMY321 (YJS5431 in this study) reference sequence allowed for single nucleotide variants detection. From 84,890 to 502,399 SNPs were detected per isolate, distributed among 811,159 polymorphic positions. A minimum of 84,500 heterozygous variants was called per sample, revealing that our dataset was devoid of haploid or homozygous isolates. The allele frequency at heterozygous sites was used to infer the ploidy of each isolate: 39 isolates showed values centered around 0.5 and were therefore considered to be diploid. The remaining 14 isolates had an enrichment of values centered around 0.33 and 0.66 and were consequently defined as triploid (Figure S1).

Interestingly, the number of homozygous variants was in the same range for all the strains in our collection with the exception of a group of 10 diploid strains. This unique group included the UMY321 reference isolate, and less than 350 homozygous variants were called per strain (Table S2). By contrast, the number of heterozygous sites was much higher in triploid isolates compared to the diploid ones (Figure S2), which is in accordance with previous studies, showing that two Australian triploid isolates were constituted of a moderately heterozygous diploid genome in combination with a divergent haploid genome (Borneman, et al. 2014; Curtin, et al. 2007).

Overall, the genetic diversity estimated by the average pairwise difference between strains is relatively high (π = 1.2 × 10^−2^) compared to *S. cerevisiae* (π = 3 × 10^−3^). As expected, the genetic diversity is lower in coding regions (π = 1.0 × 10^−2^) compared to intergenic regions (π = 1.5 × 10^−2^). In addition, SNPs found in CDS (CoDing Sequence) are mostly synonymous (64%) while nonsense mutations are rare (1%). These latest mutations also show a lower allele frequency within the population and a more contrasted but similar pattern for observed for missense variants (Figure S3).

### Phylogeny and strain relatedness in *B. bruxellensis*

The 811,159 polymorphic positions were used to infer phylogenetic relationships between the isolates. To handle heterozygosity within our samples, heterozygous SNPs were encoded using the IUPAC code and the average state method was used for the distance calculation. Five distinct clusters were highlighted in the neighbor-joining tree, three of them (G2N1-3) were composed of diploid isolates while the remaining two (G3N1-2) exclusively contained triploid isolates (Figure 1).

**Figure 1.**
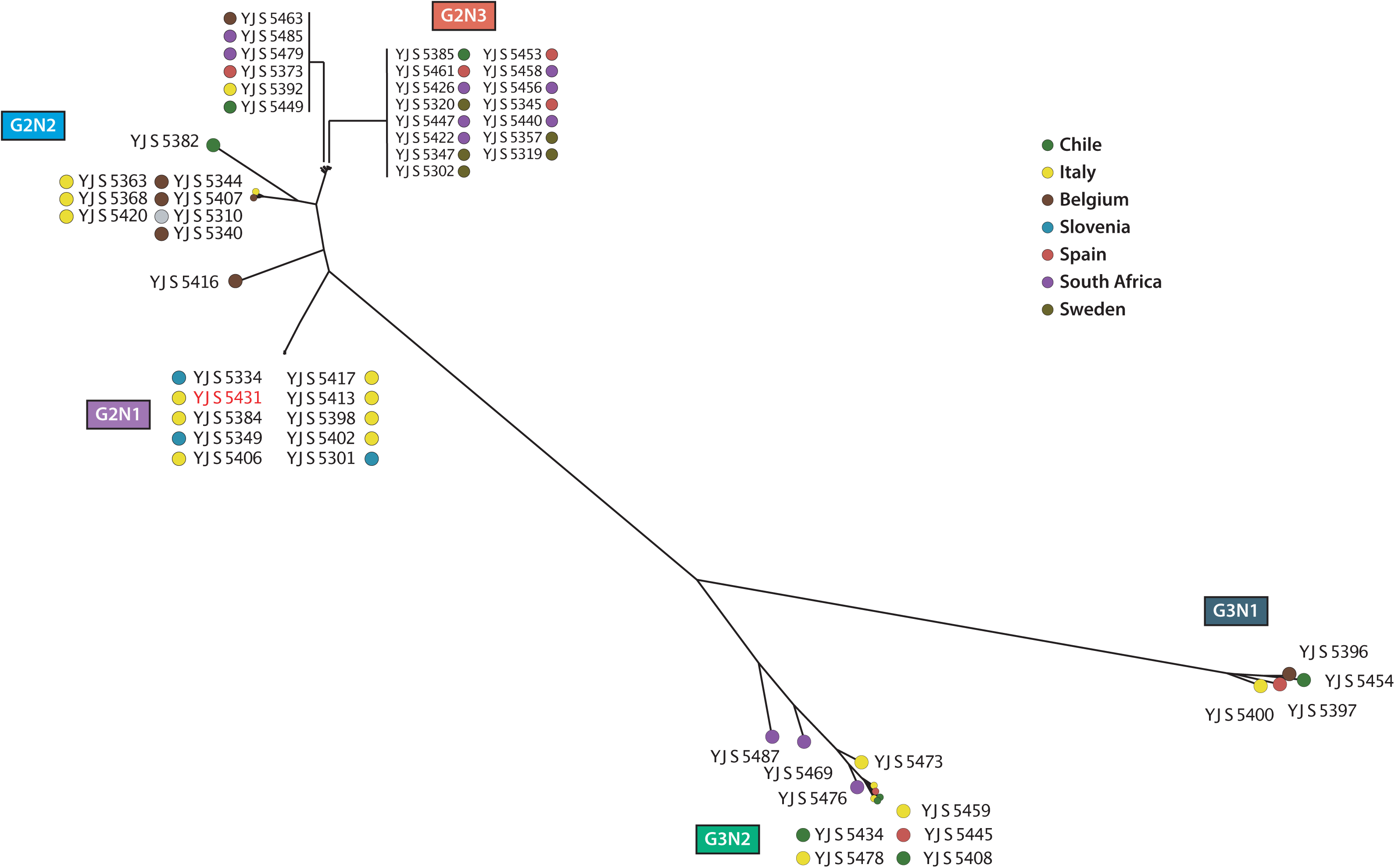
Neighbor-joining tree of the 53 isolates based on the 811,159 detected polymorphic positions. We identified five clusters composed of diploid (G2) or triploid (G3) strains. YJS5382 is the only triploid strain found among the diploid clusters (left side).

These clusters are mostly in accordance with the five major clusters recently described using microsatellites on a very large population (Avramova, et al. 2018). For example, the two triploid clusters, G3N1 and G3N2, correspond respectively to the so-called triploid beer AWRI1608-like cluster and the triploid wine AWRI1499-like cluster. Moreover, the three distinct diploid clusters can be observed, enhancing the delineation of this population. The G2N3 cluster corresponds to the wine CBS2499-like cluster, while the G2N1 cluster (including the wine reference YJS5341 strain isolated in Italy) is distinct, although both were in a single Kombucha-like cluster in the former analysis.

The YJS5382 isolate could not be assigned to any cluster in our analysis, which is also in accordance with this previous study (Avramova, et al. 2018), as it is the only representative of the wine L0308-like cluster in our dataset. Interestingly, this isolate is the only triploid that does not group with the G3 clusters. By contrast, it is in fact closer to the diploid clusters, suggesting an independent triploidization event for this strain. Additionally, the YJS5416 strain is not closely related to any cluster in our tree and is located between the G2N1 and G2N2 clusters and it might, therefore, be a representative of a new subpopulation.

### Genomes are punctuated by a few loss-of-heterozygosity regions

To have a better insight into the nucleotide variation along the genome, we examined the genetic diversity across non-overlapping windows of 10 kb for each cluster (Figure 2). Overall, the triploid subpopulations show a higher genetic diversity (π_G3N1_ and π_G3N2_ = 2 × 10^−2^) compared to diploid clusters (π_G2N1_ = 3.5 × 10^−3^, π_G2N2_ = 6 × 10^−3^, π_G2N3_ = 4 × 10^−3^). However, the nucleotide diversity is not homogeneous along the genome (Figure 2). In fact, the two triploid clusters display a lower genetic diversity (π < 1 × 10^−2^) within a large region of ~ 1.2 Mb on the left side of the scaffold 1. Interestingly, two similar regions can be observed in the triploid wine subpopulation (G3N1) on the right extremity of the scaffolds 3 and 5. In the diploid subpopulations, similar regions exhibiting a low genetic diversity can also be observed on almost all chromosomes. In the G2N2 and G2N3 clusters for which a notable decrease of the Tajima’s D values is also observed, indicating a high proportion of rare alleles in these regions compared to the rest of the genome. While the Tajima’s D value never drops below 0 with the exception of the right side of the scaffold 6 in the G2N1 cluster, the impacted region showed a lower Tajima’s D value. This difference might be due to the close proximity between the strains belonging to this cluster and the reference sequence.

**Figure 2.**
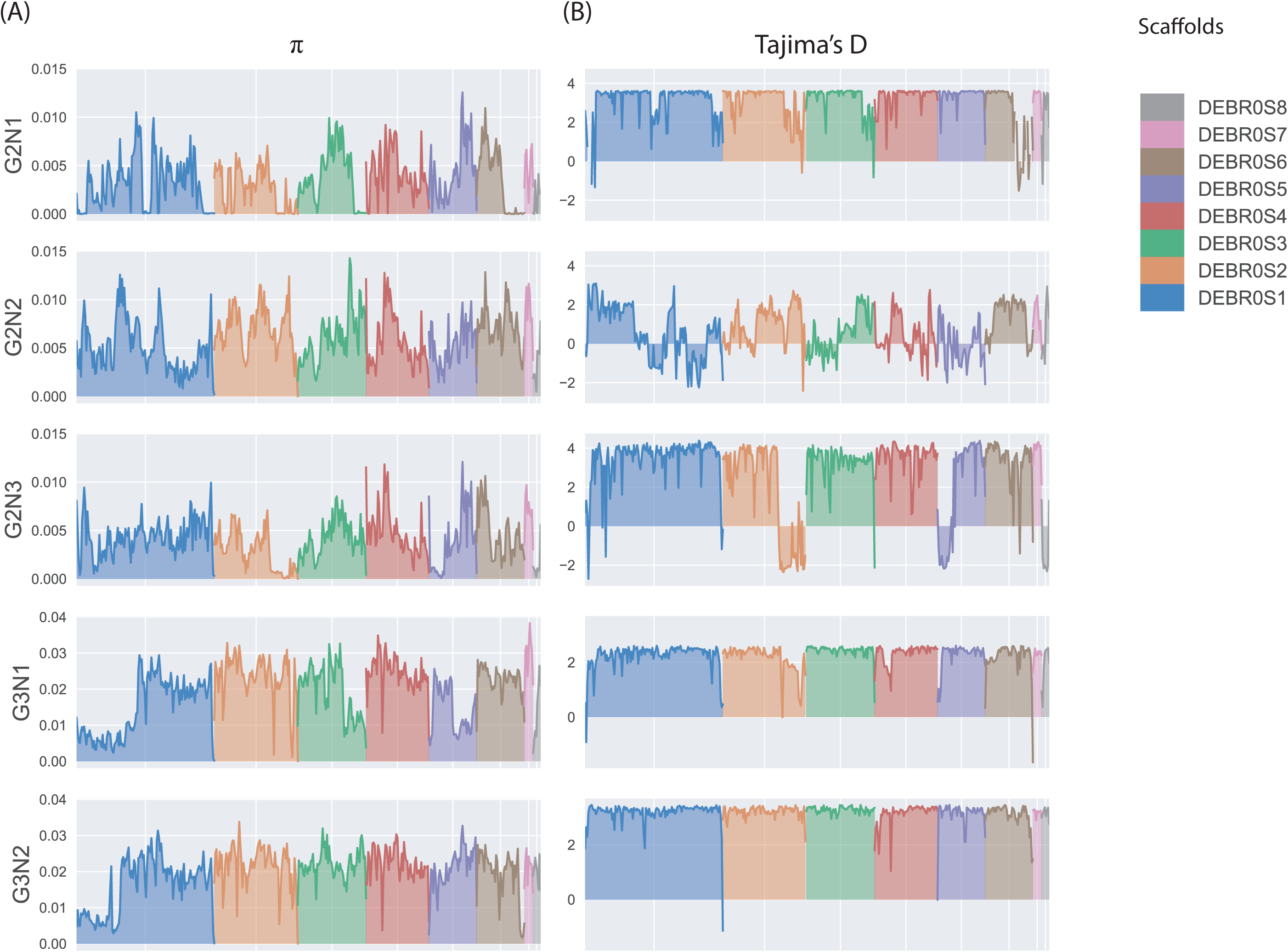
Variation of nucleotide diversity metrics for each cluster using non-overlapping window (10 kb). (A) Pairwise nucleotide diversity (π), (B) Tajima’s D values.

To better understand the origins of these regions with lower genetic diversity, we scanned the genomes for the presence of loss of heterozygosity events (LOH). This type of event leads to a decrease of the genetic variability allowing the expression of recessive alleles, and therefore can result in a beneficial adaptation in some environments. The prevalence of LOH events has recently been observed in several yeast species such as *S. cerevisiae* (Peter, et al. 2018), *Candida albicans* (Ropars, et al. 2018) and *Kluyveromyces marxianus* (Ortiz-Merino, et al. 2018). Moreover, previous studies of *B. bruxellensis* isolates revealed the presence of several regions which underwent LOH. This was particularly clear in the case of the wine strains AWRI1613 and CBS2499 for which 17.9% and 16.3% of the genome are impacted, respectively (Borneman, et al. 2014). To detect LOH regions in our collection, the allele frequency of the polymorphic sites was plotted along their chromosomal location in the reference assembly and the number of heterozygous SNPs along the genome was further examined using 10 kb-sliding windows (Figure S4).

In all diploid isolates, several regions of LOH were observed representing a total of approximately 0.7 to 2.7 Mb (mean value of 1.8 Mb, 13% of the genome) distributed among 6 to 18 regions (mean value of 11.7 regions) (Table S3). This level of LOH is low compared to *S. cerevisiae* for which LOH cover approximately 50% of the genome on average (Peter, et al. 2018). The patterns of LOH are very well conserved within clusters and only very few regions are strain-specific. Some LOH events seem to have arisen before cluster expansion as they are shared between several clusters. Moreover, most of the identified regions also showed a low nucleotide diversity, suggesting that LOH is the main reason for the nucleotide variability patterns mentioned above. In triploid isolates, regions exhibiting a complete loss of heterozygosity are uncommon and only 1 to 3 LOH regions were detected covering 29 to 279 kb. These regions are well conserved between the strains. Interestingly, several regions in the triploid strains showed a reduced heterozygous rate and a lower nucleotide diversity. However, they did not meet our criteria to be considered as LOH. These regions could have resulted from an ancient LOH event that occurred before the separation of the two triploid clusters and the accumulation of new mutations, which is especially possible for the left part of the scaffold 1.

Masking LOH regions allowed us to precisely determine the level of heterozygosity, which ranged from 6.2 to 36.3 heterozygous sites per kb (Table S4). As expected, all diploid isolates share the same level of heterozygosity, with a mean value of 7.1. Much higher heterozygosity rates were associated with triploid strains: YJS5382 has 11.4 heterozygous SNPs/kb, and this level is much higher within the triploid clusters. Indeed, a mean of 34 heterozygous SNPs/kb was observed, this level was slightly higher in the G3N2 cluster compared to the other triploid subpopulations with 31 and 35 heterozygous SNPs/kb, respectively. This difference is most likely due to the regions that have undergone LOH in the G3N1 cluster and are not found in the G3N2 strains.

### Aneuploidies and segmental copy variants are uncommon in *B. bruxellensis*

To determine the frequency of aneuploidy as well as of segmental copy variants in our collection, we examined the coverage distribution along the reference genome using non-overlapping windows of 20 kb. Coverage deviations were confirmed with the allele frequency variation in these regions. We observed such deviations at the whole scaffold level for only 3 isolates (5.6% of the 53 strains), and additionally only related to the smallest scaffolds (scaffolds 7 and 8). This result suggests that aneuploidies are rare in *B. bruxellensis* compared to other species such as *S. cerevisiae*, for which analysis of 1,011 strains revealed that roughly 20% of the population is affected by aneuploidies (Peter, et al. 2018). Segmental variations were more prevalent and detected in 17 isolates (Figure 3), 9 of which harboured several regions that were affected. However, the relatively weak prevalence of these events and the size of the affected regions (mean size approximately 350 kb) suggest that they could not explain the highly variable karyotypes observed within this species alone and that balanced rearrangements may also occur. Moreover, triploid clusters display a higher proportion of samples carrying segmental variants with more than half of the samples affected by such variants (Figure S5). This result supports the idea that hybrids, and generally polyploid strains are affected by structural changes (Otto, et al. 2007).

**Figure 3.**
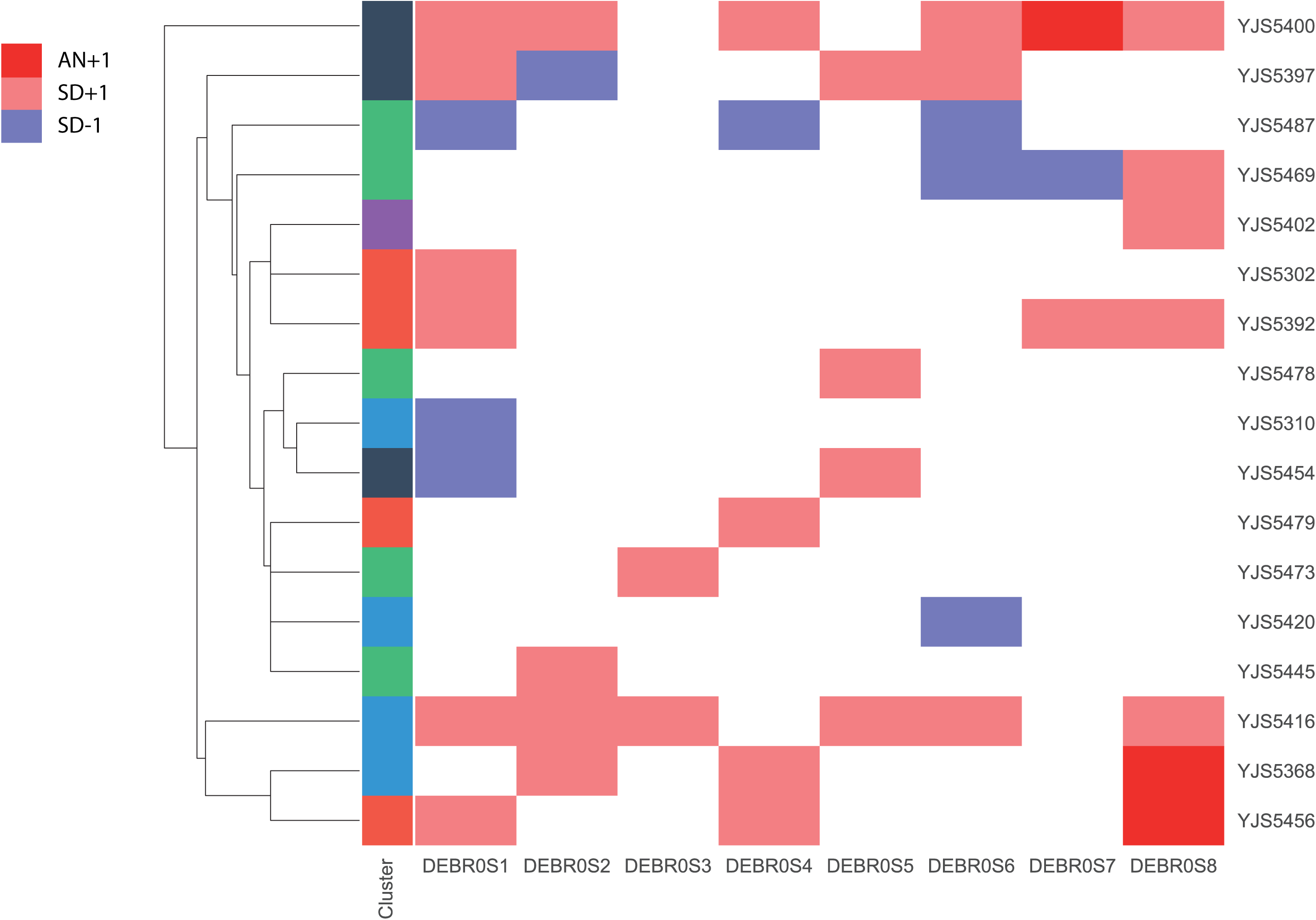
Distribution of aneuploidies (AN) and segmental duplications (SD) for each sample.

We then focused on triploid isolates to determine whether these copy variants affected the haploid or diploid genomic version by taking advantage of the allele frequency within these regions. Indeed, while a duplication of the haploid version will result in an equal genome version ratio (2:2) and therefore in a frequency shift to 0.5, a supplemental copy of the diploid genome will lead to a frequency of 0.25 and 0.75 (3:1). On the other hand, a deletion of the haploid and diploid versions will result in an allele frequency of 1 (2:0) and 0.5 (1:1) in these regions, respectively. Eight triploid strains carrying segmental duplications were investigated (Figure S6, Table S5). A total of seven duplicated regions showed a 3:1 ratio *vs* a 2:2 ratio, suggesting that duplication of the diploid genome is more common. Moreover, six out of the seven deleted regions showed a ratio close to 0.5, corresponding to the loss of one copy of the diploid genome. This result suggests that the haploid version of the *B. bruxellensis* hybrids is mostly conserved compared to the diploid, in regard to deleterious structural variants.

### Triploid genomes are more subject to gene copy number variation than diploids

Our sequencing data provided the opportunity for a more precise view on the variation of gene copy numbers (CN) in the five groups of *B. bruxellensis*. By scanning the coverage for each strain using a GC content normalized approach (see Methods), we detected the genes impacted by a variation of the CN. Among the 5,226 annotated protein-coding genes in the reference genome, 4,088 genes (78.2%) showed a variation in the number of copies in at least one strain. Most of these genes are present as duplicates (N = 3,587). Deleted copies were found for 1,734 genes, among which only 100 were totally absent from the genomes. However, gene deletions are more shared across isolates compared to duplications (mean of 4.70 *vs* 2.72, respectively). In addition, CNs are enriched in subtelomeric regions as previously observed in other yeast species such as *S. cerevisiae* (Peter, et al. 2018), and LTR, mobile elements and tRNA are proportionally more impacted by these variants compared to CDS (Figure S7).

Interestingly, CDSs in triploid isolates are more subjected to CN variation (Figure 4A, p-value = 2.17 × 10^−128^), which is consistent with the segmental duplication profiles. While the number of CNVs in diploid strains is 5.6 times lower compared to triploid strains (mean of 169 *vs* 881, respectively), this value is variable among strains and some diploid isolates, such as YJS5456, display a number of CNVs similar to that observed in triploid strains (Figure 4B, Table S6). Moreover, the ratio between deleted and duplicated CDS is variable among clusters (Table S6) and the G3N1 cluster shows a significantly higher number of duplicated CDSs compared to those that have been deleted (ratio = 2.6, Table S6). This could obviously be linked to the high number of segmental duplications found in the G3N1 cluster.

**Figure 4.**
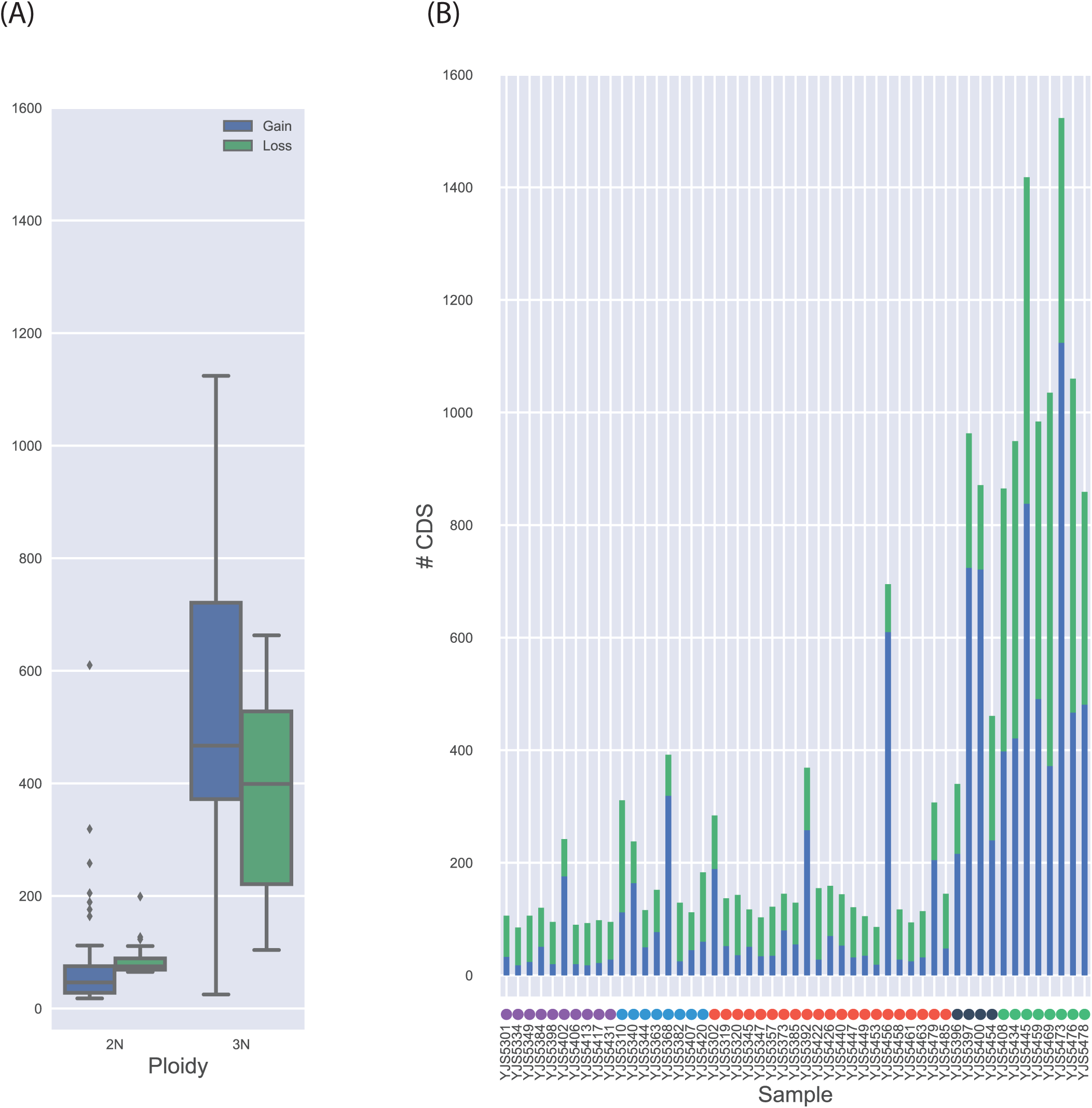
CNV distribution among the population. (A) Distribution of CNVs by ploidy. (B) Number of CNVs for each sample.

### Functional insight into the duplicated and deleted genes

Gene copy variants are known to be a driving mechanism of genomic adaptation to changing environments in yeast (Kondrashov, et al. 2012) and are frequently found to be associated with domesticated processes. For example, in *S. cerevisiae*, duplications of the *CUP* genes have been repeatedly associated with resistance to copper (Peter, et al. 2018, Strope, et al. 2015) and the cluster of *MAL* duplicated genes has been highlighted as facilitating the utilization of maltose. It can be found in the ale beer isolates for which this sugar is the main carbon source during the fermentation process (Gallone, et al. 2016). Within *B. bruxellensis*, the investigation of the genome assemblies related to different isolates already highlighted cases of gene content variation between strains. For example, an expansion of the alcohol dehydrogenase family, enabling greater control over ethanol formation and consumption, has been found in the wine isolate AWRI1499 (Curtin, et al. 2012). In another study, comparisons of two wine and one beer strains highlighted 20 genes encoding sugar metabolic processes and nitrogen consumption. These genes were found to be present in the wine strains AWRI1499 and CBS2499 but missing in the beer strain ST05.12/22 (Crauwels, et al. 2014).

To determine whether these variations are shared between samples or are independent events, we first investigated the gene copy numbers in the whole population using the 20 candidate genes identified in a previous study (Figure 5). We found that all these genes were only missing in the YJS5392 strain (G2N3). This strain was isolated in Belgium and is closely related to the STO5.12/22 beer isolate. However, this strain is an exception within the whole population and most of the genes are present in the different subpopulations. Interestingly, two genes coding for MFS drug transporters and one gene coding for a high-affinity glucose transporter have more than four copies in all triploid strains. Moreover, some strains from the triploid cluster G3N2 related to the wine strain AWRI1499 have multiple copies of genes coding for proteins responsible for galactose, glucose, and hexose transporter and metabolism while diploid strains have two copies, and one copy is found in isolates from the G2N1 cluster. Finally, genes involved in nitrogen metabolism were completely absent from the seven diploid strains independently found in the three different clusters.

**Figure 5.**
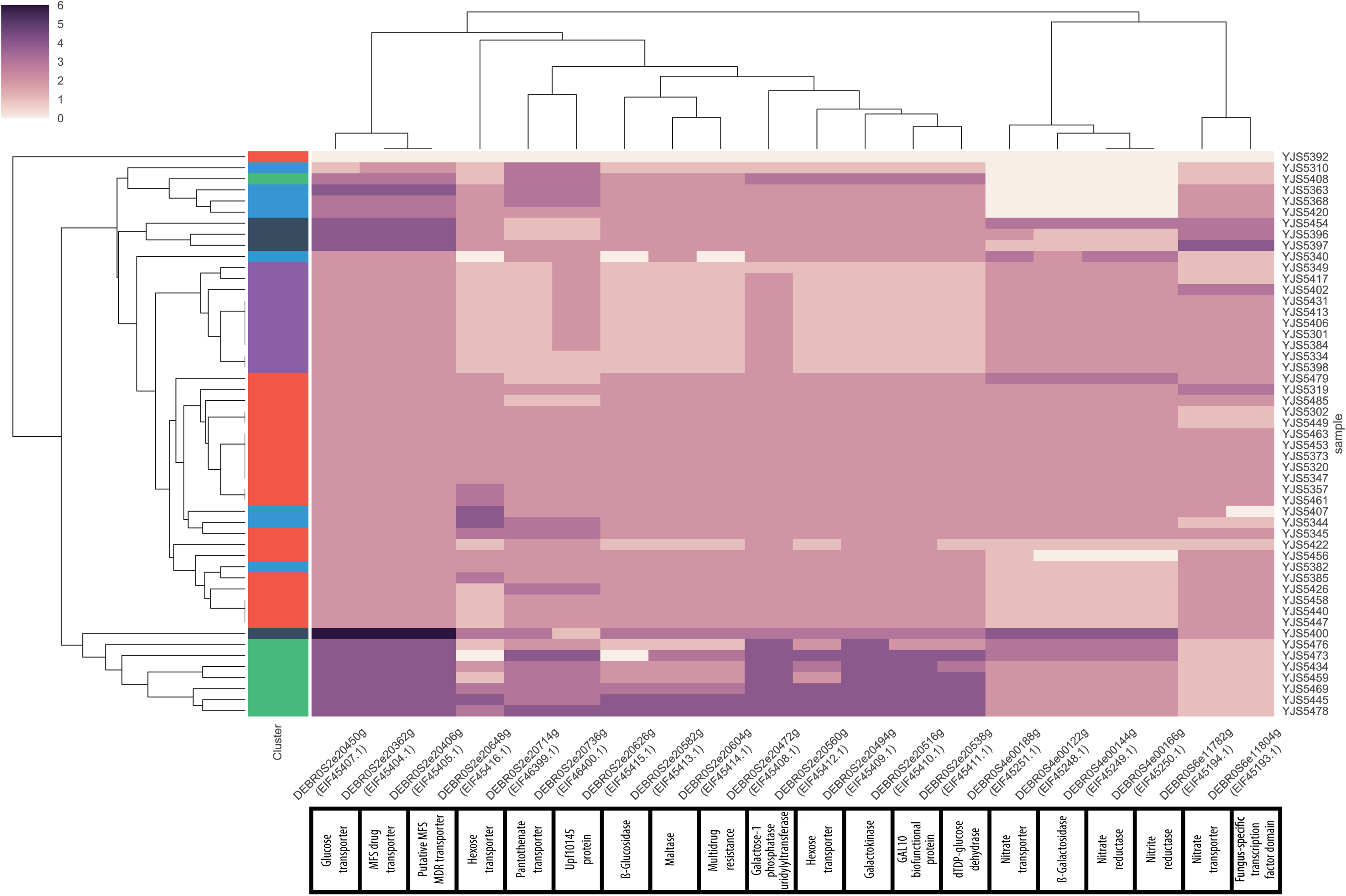
CNVs within the whole population for 20 genes previously found to be present in the two strains AWRI1499 and CBS2499 but missing in the ST05.12/22a isolate.

To determine if other genes are under selective pressure within a whole subpopulation, we examined all duplicated or deleted genes within each cluster for which at least 70% of the strains displayed the same structural changes (Table S7). Most of these genes were found in the triploid clusters, especially in the G3N2 cluster encompassing strains related to the AWRI1499 wine strain. Several genes involved in core pathway, such as histones H3 and H4 or ribosomal proteins can be found in the G2N3 and G3N2 clusters. Moreover, GO-Term analysis of genes found in the G3N2 cluster revealed enrichment for several biological processes including galactose metabolism (*GAL7*, *GAL3*, p-value = 0.003), nitrogen utilization (*UGA1*, *ATO2*, p-value= 0.004) and transmembrane transport (9 genes, p-value = 0.005).

### The *B.* bruxellensis pangenome is small with a few accessory genes

To complete our analysis of the gene content within *B. bruxellensis*, we determined the species pangenome, *i.e*. the global set of ORFs (Open Reading Frame) present within the species. To that end, *de novo* assemblies for all the studied isolates were constructed and scanned to detect non-reference materials (see Methods). In fact, 203 (3.7%) additional protein-coding genes were detected, leading to the identification of a total of 303 accessory genes (5.5%), as 100 genes were found as fully deleted. This result shows that the pangenome is much smaller compared to that defined in *S. cerevisiae*, with a total of 1,712 accessory genes (Peter, et al. 2018). In *B. bruxellensis*, 4,923 genes were found in all isolates and consequently were assigned to the core-genome (94.5% of the pangenome). Supplemental ORFs were mostly found in triploid strains, with 154 and 49 of ORFs specific to the triploid and diploid strains, respectively. A significant part of these supplemental ORFs can be found in only one isolate (33%). However clustering analysis still revealed groups of genes associated with the different subpopulations (Figure S8). To determine if these genes provide adaptive advantages, we searched for putative functions based on similarity searches from the protein sequences. Several transporters were found to be specifically shared within the triploid wine related cluster (G3N2), such as two accessory genes with similarities to MFS drug transporters. However, no significant functional enrichment was found and multiple genes were linked to transposons or flocculation proteins, which are known to easily degenerate during evolution and could therefore be false positives in our analysis. While current data does not provide the opportunity to determine the precise origin of supplemental genes, their prevalence in the triploid isolates suggests that a significant number of these genes may result from hybridization events. Indeed, it is possible that the formation and selection of hybrids is linked to the contribution of the two genomes, sharing both new alleles and genes in a single individuals, ultimately conferring beneficial advantages to specific environments. In this regard, further studies based on haplotype phasing in hybrid genomes need to be initiated, and will provide valuable insights into the genomic adaptations driven by hybridization events.

## Conclusion

With the advent of affordable sequencing technologies, it is now possible to explore and analyse the genome-wide variability within non-model, but industrially relevant species. In this study, for the first time we provide a comprehensive description of the genetic variability at the genome-scale of a population of *B. bruxellensis*, giving us a better view of its evolutionary history and the genetic variations underlying the phenotypic diversity within this species. Our results show the presence of at least two hybridization events, which is one of the main factors involved in the division classification of this species into subpopulations. It is likely that they are a driving mechanism of *B. bruxellensis* evolution as an adaptive response to the harsh environments found in the domestication processes. Interestingly, similar and common patterns of genetic variability are observed in both wine and beer triploid subpopulations. Nevertheless, significant differences between the genomes of the two subpopulations can also be found, suggesting the presence of potential genomic adaptations, which are specific to each of them. In addition, our results indicate that loss of heterozygosity is also present in *B. bruxellensis* evolution, impacting the genetic diversity within the species. Moreover, several aneuploidies, segmental duplications, and CNVs were also found in the whole population, especially in the triploid strains, indicating that these genome dynamics are important within the species and that triploid hybrids favour these such types of structural variations. These observations are similar to what has been shown in the hybrid species *S. pastorianus* resulting from the combination of the genomes of *S. cerevisiae* and *S. eubayanus*, for which subpopulations show extensive chromosome loss and LOH events (Okuno, et al. 2016). However, whether the hybridization events in *B. bruxellensis* derived from isolates of the same or two different species remains unknown and both a deeper analysis of triploid isolates genomes and the sequencing of closely related species genomes would be needed to investigate this aspect of their biology. At the whole population scale, our data provides the opportunity for a deeper view of the genetic variants involved in the phenotypic diversity of *B. bruxellensis*. Analysis of CNVs and accessory genes in the populations highlighted several genes involved in drug and sugar transports as previously found in other analyses. However, these genes were mostly found in the wine related triploid clusters. Interestingly, the nitrogen pathway was independently lost in several diploid isolates within different subpopulations, suggesting that nitrate assimilation is not a common requirement for *B. bruxellensis* isolates. This result is in accordance with a phenotypic analysis in which up to a third of the isolates failed to grow on nitrate, which could result from the reduction of ethanol and the production of acetic acid during anaerobic fermentation nitrate assimilation (Galafassi, et al. 2013).

## Materials and methods

### Isolates and sequencing

This study was performed using a collection of 53 *B. bruxellensis* isolates originating from diverse ecological and geographical origins (Table S1). Most of the samples were isolated from Europe, but some also originated from South Africa, Australia and Chile. While a subset of the samples had no ecological origins associated with them, approximately half of them were retrieved from fermentative and wine related environments.

All *B. bruxellensis* isolates were subjected to Illumina paired-end sequencing. Yeast cell cultures were grown overnight at 30°C in 20 mL of YPD medium to early stationary phase before cells were harvested by centrifugation. Total genomic DNA was then extracted using the QIAGEN Genomic-tip 100/G according to the manufacturer’s instructions. Genomic Illumina sequencing libraries were prepared with a mean insert size of 280 bp and subjected to paired-end sequencing (2 x 100 bp) on Illumina HiSeq 2500 sequencers.

### Reference sequence correction and annotations

The annotation of the UMY321 genome sequence (Fournier, et al. 2017) was performed using Amadea Annotation transfer tool (Isoft, France) with *Lachancea kluyveri* CBS3082^T^ and *Debaryomyces hansenii* CBS767^T^ genomes as references (revised versions available at http://gryc.inra.fr). This step was followed by a manual curation with RNA-Seq data from *Brettanomyces bruxellensis* CBS2499 (Sequence Read Archive SRR427169^17^). Tophat2 aligner tool v. 2.1.0 was used to map reads against the assembled genome of UMY321 (Kim, et al. 2013). Artemis v16.0.0 was used to visualize BAM files, to adjust exons and introns coordinates and to identify lncRNA (Carver, et al. 2012). tRNA genes were detected using tRNAscan-SE v1.3.1 (Lowe, et al. 2016). Transposable elements were identified by BLAST with known yeast elements from different families such as *Ty1-copia*, *Ty3-gypsy*, and *hAT*, as queries.

A high number of out-of-frame genes were predicted, mostly related to long read sequencing errors. To correct these sequencing errors, an independent assembly was constructed by running SOAPdenovo from Illumina short reads (Luo, et al. 2016). This latter was scaffolded with Redundans (Pryszcz, et al. 2016) using the initial UMY321 assembly (Fournier, et al. 2017) as template. The sequences corresponding to the frameshifted genes were detected in the new assembly by similarity searches and inferred in the original assembly if the correct frame was recovered. The subsequent annotations were modified accordingly in the UMY321 assembly.

### Mapping, variant calling and annotation

For each sample, Illumina paired-end reads were mapped on the corrected sequence of UMY321 genome with BWA mem (v0.7.12, default parameters) (Li and Durbin 2009). Alignments were post-processed using SAMtools fixmate (v1.1) (Li, et al. 2009), GATK realignment (v3.3.0) (McKenna, et al. 2010) and Picard MarkDuplicates (v1.1.140) (broadinstitute.github.io/picard).

Aneuploidies and segmental duplication events were visually identified based on read coverage profiles. To that end, coverage was computed for each position with SAMtools depth function and mean coverage values along the genome were plotted using 20 kb non-overlapping sliding windows. Single nucleotide polymorphisms and small indels were called using GATK Haplotype Caller function with a ploidy set to 2. In order to determine the frequency of each variant for ploidy analysis, VCF file was annotated using GATK Variant Annotator function. Variants with a coverage lower than 10 X or a quality score lower than 25 were removed using bcftools (v 1.5) filter (Li, et al. 2009). Finally, variant files were merged into one single file using vcftools (v. 0.1.13) merge function (Danecek, et al. 2011). To assess the impact of variants on protein sequences, annotation was processed with SnpEff (v 4.3i) (Cingolani, et al. 2012) and the impact on protein function for non-synonymous SNPs was predicted using SIFT (v 6.2.1) (Kumar, et al. 2009).

### Allele frequency and nucleotidic diversity

The allele frequency of the heterozygous single nucleotide variants was determined using the ABHet annotations generated by GATK Variant Annotator.

To get an estimate of the scaled mutation rate, the average pairwise nucleotide diversity (π), the proportion of segregating sites (θ_w_) and Tajima’s D value were computed with variscan (v. 2.064) (Tajima 1989; Vilella, et al. 2005). It was run with mode 12, usemut 1 and 10-kb non-overlapping windows on multiple alignments of the concatenated chromosomes with SNPs inferred from each sample. A similar process was used to determine these estimators for coding and non-coding regions.

### Phylogenetic relationships

The phylogenetic relationships between samples were estimated through the construction of a neighbor joining tree based on the BioNJ algorithm provided by SplitsTree4 (Huson, et al. 2009). To that end, a multiple alignment of the concatenated contigs was produced in which SNPs were inferred using the IUPAC code, leading to a single sequence for each sample. The MatchState option was selected to properly manage the heterozygous positions while calculating the distances.

### LOH detection

The genome of each isolate was scanned (50 kb sliding windows with 25 kb overlap) to identify regions with 10 or less heterozygous sites per 50 kb that were considered to be under loss of heterozygozity (LOH).

### Copy number variants

Control-FREEC (v10.6) (Boeva, et al. 2011) was used to determine the number of copy of 1 kb region along the genome of each sample, with the following parameters: breakPointThreshold = 0.6, window = 1000, telocentromeric = 6000, step = 200, minExpected GC = 0.3 and maxExpectedGC = 0.5. The number of copy of each gene was deduced from these data and normalized according to the ploidy of the isolate, allowing for the detection of the copy number variants.

### Supplemental genes identification

To determine the set of supplemental genes (genes not found in the reference genome) within our population, we used a pipeline similar to the one developed for the analysis of 1,011 *S. cerevisiae* strains (Peter, et al. 2018). We first constructed an assembly for each sample using Abyss (v2.0.2) with 67 as kmer size (Jackman, et al. 2017). Assemblies were compared to the reference sequence with BLASTn allowing for the determination of non-reference segments. Gene prediction tools were applied on these segments through both SNAP (v2006.07.28) (Korf, et al. 2004) and Augustus (v. 3.2) (Stanke and Waack 2003) with training sets generated using the *B. bruxellensis* proteome. Genes showing an excess of low complexity region as well as those whose product had a length less than 50 amino acids were removed. All genes were then compared to each other with BLASTp and the application of a graph-based method leads to the identification of a non-redundant set of supplemental genes for the whole species.

Finally, these genes were annotated through similarity searches against the reference proteome and several protein databases such as Uniref90, SwissProt Fungi, UniProt Fungi as well as Saccharomycotina species found in the SSS (http://sss.genetics.wisc.edu/cgi-bin/s3.cgi) and GRYC (http://gryc.inra.fr/) database.

## Data availability

The sequences related to the YJS5363, YJS5431 and YJS5447 were previously released in the European Nucleotide Archive under the accession number PRJEB21262 (Fournier, et al. 2017). All sequencing data generated for this study have been deposited under the accession number PRJEB28420. The annotations of the UMY321 isolate were deposited under PRJEB33245.

## Acknowledgments

We are grateful to Warren Albertin and Isabelle Masneuf-Pomarede for fruitful discussions and invaluable advice. We thank the BioImage platform (IBMP-CNRS, Strasbourg, France) for their support. This work was supported by the Agence Nationale de la Recherche (ANR-18-CE20-0003-02 and ANR-18-CE12-0013-02). J.-S.G. was supported by a grant from the French “Ministère de l’Enseignement Supérieur et de la Recherche.” J.S. is a Fellow of the University of Strasbourg Institute for Advanced Study (USIAS) and a member of the Institut Universitaire de France.

