## Supplemental figures for "High complexity and degree of genetic variation in *Brettanomyces bruxellensis* population"

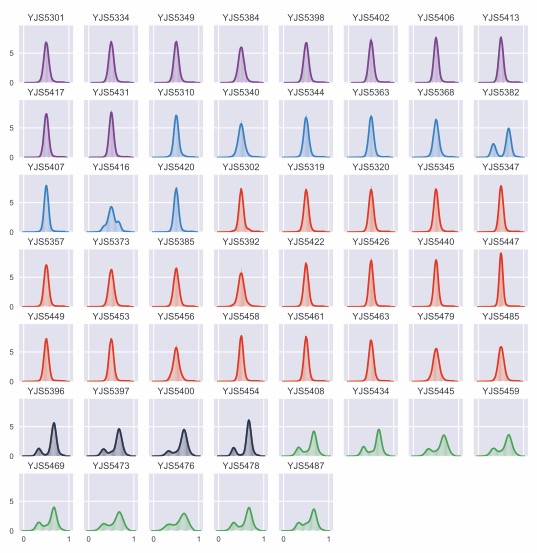

**Figure S1**. Allele frequency distribution of the heterozygous SNPs within each sample. Colors correspond to the clusters defined in Figure 1.

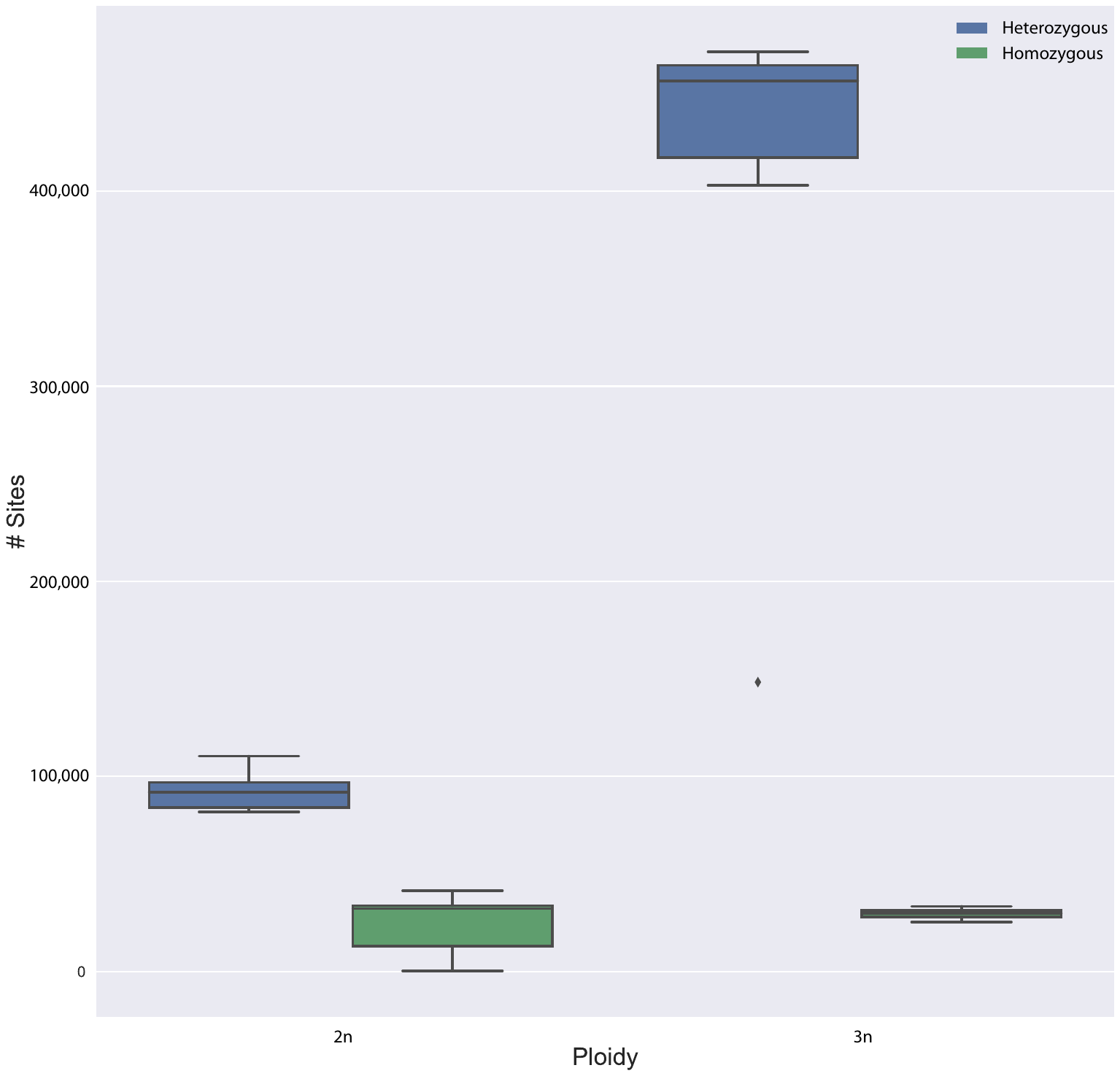
 **Figure S2**. Distribution of the polymorphic sites for each strain by ploidy.

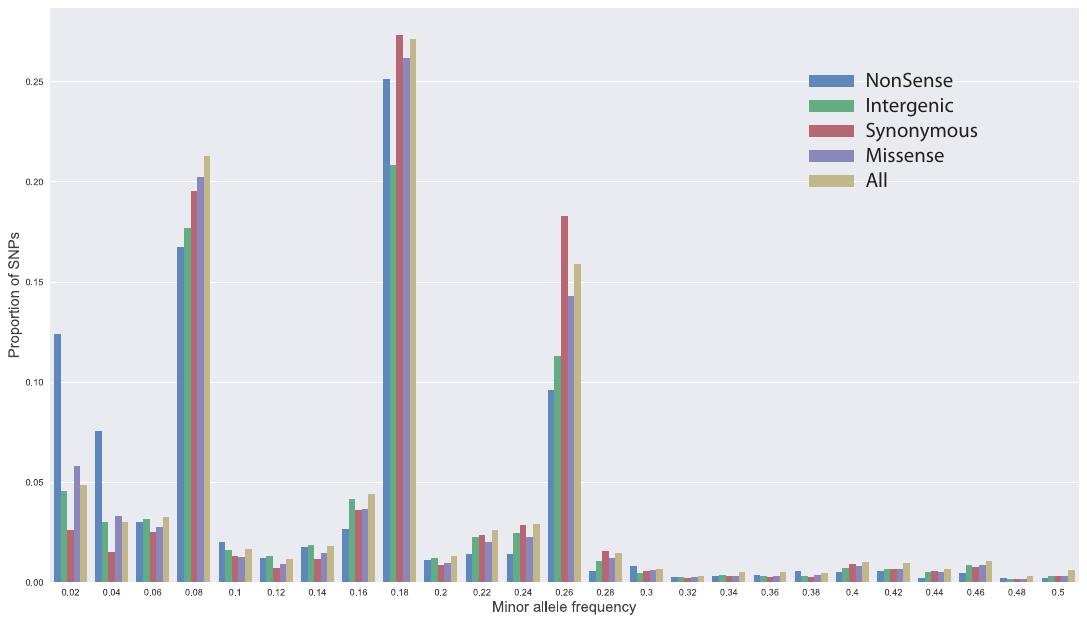

**Figure S3**. Minor allele frequency of single nucleotide polymorphism (SNP) grouped by genomic locations and functional annotation.

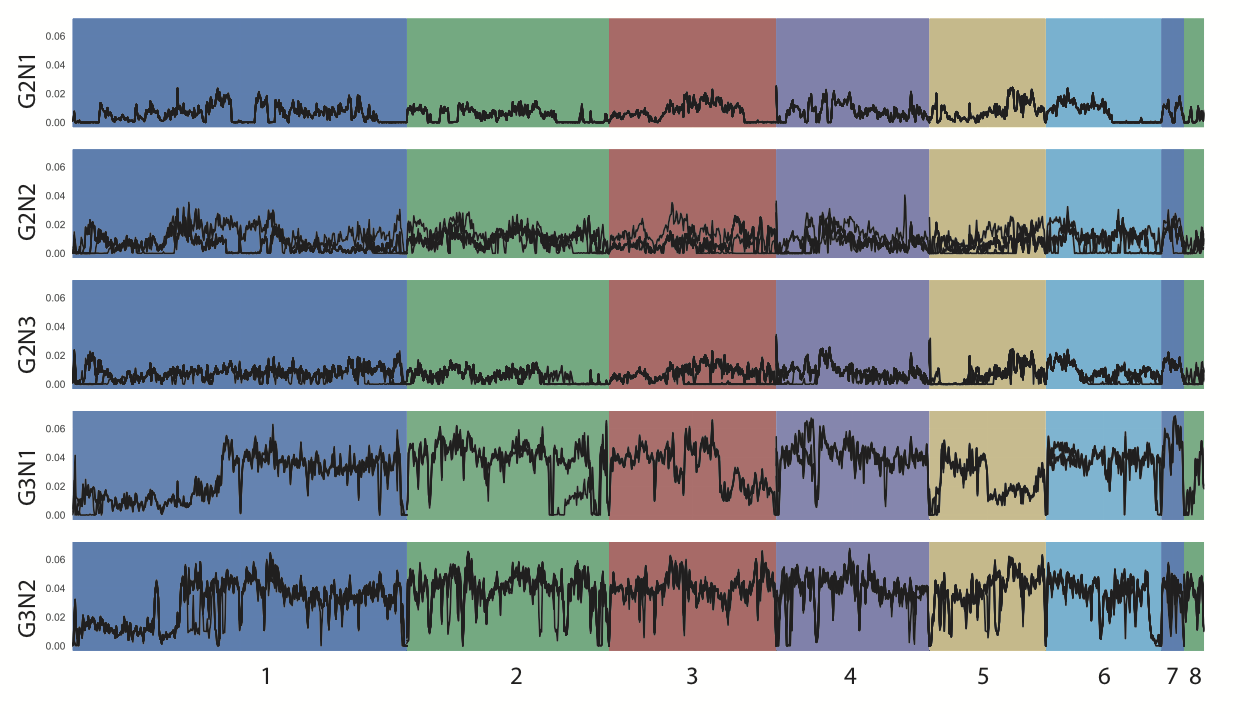

**Figure S4**. Heterozygosity level along the genome for each cluster using 10 kb-sliding windows.

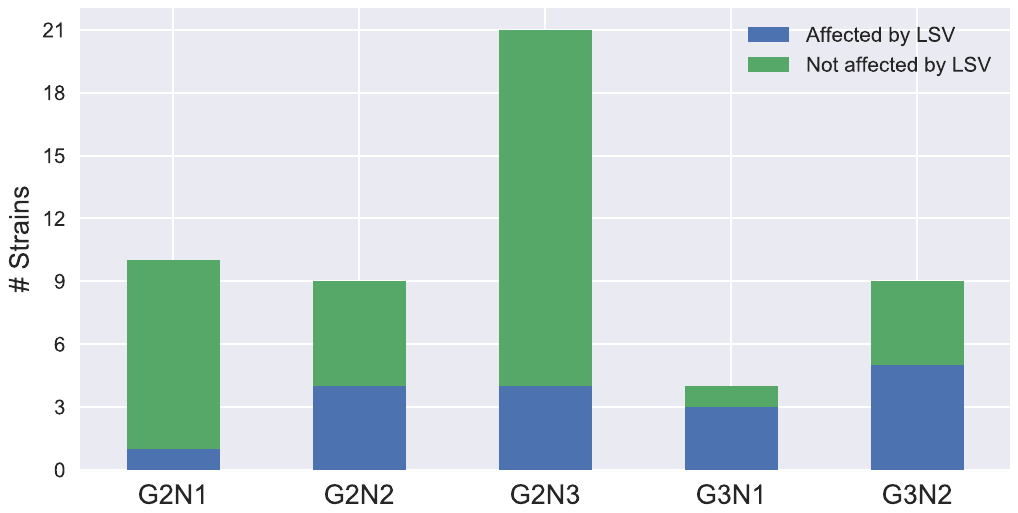

**Figure S5**. Number of strains affected by aneuploidies or segmental duplications by cluster.

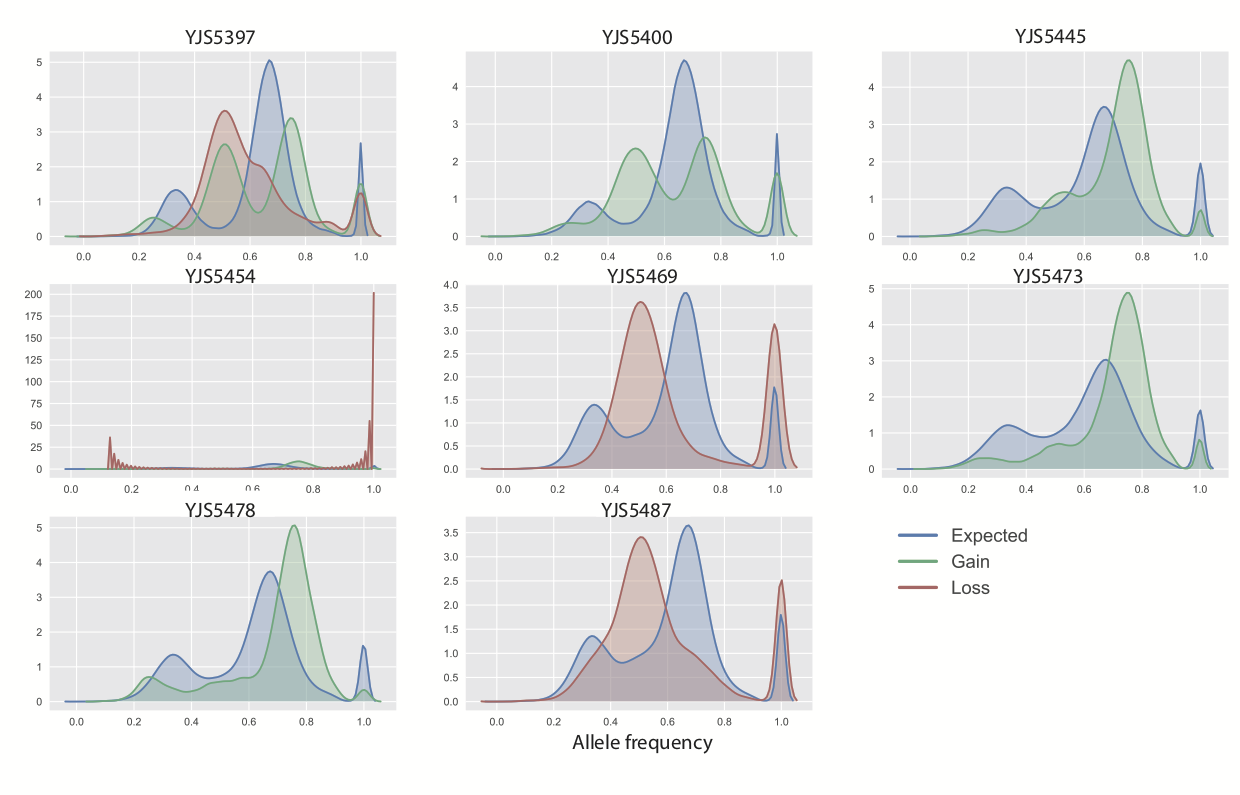

**Figure S6**. Allele frequency distributions of heterozygous SNPs within large regions that show variation in the number of copy, in triploid strains.

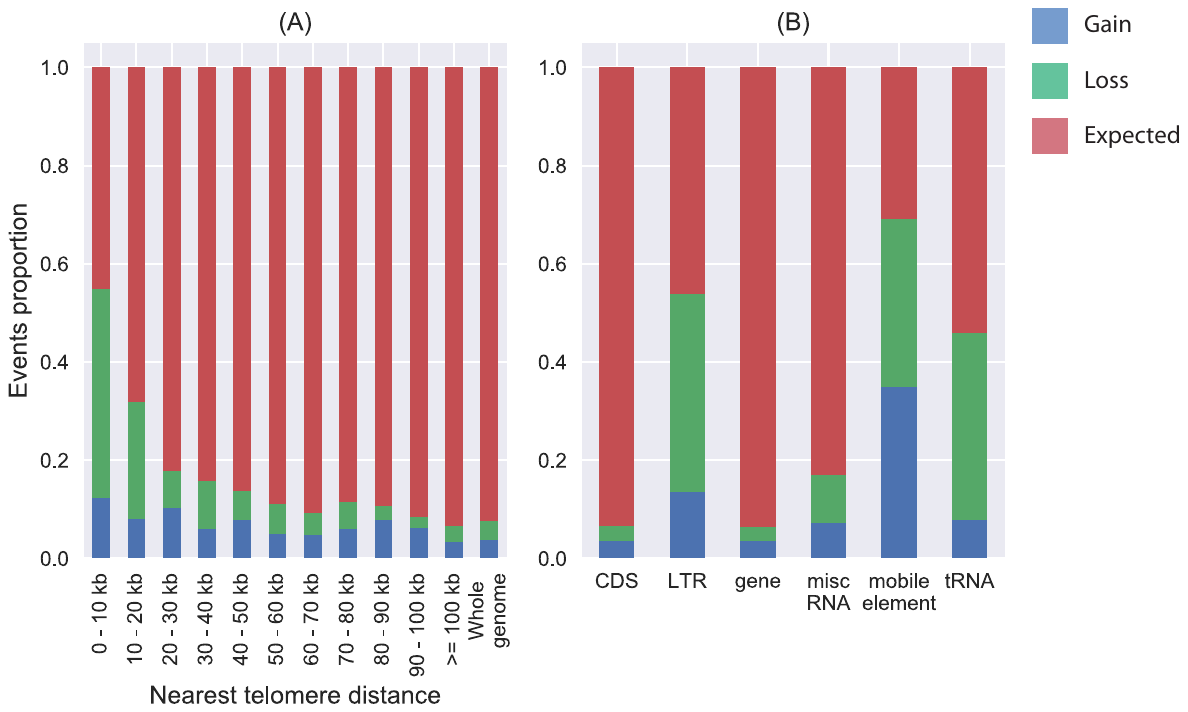

**Figure S7**. CNV distribution in the genome. (A) Fraction of CNVs affecting all genetic elements within the whole population based on their distance from the nearest telomere. (B) Proportion of features affected by CNVs within the whole population, grouped by feature type.

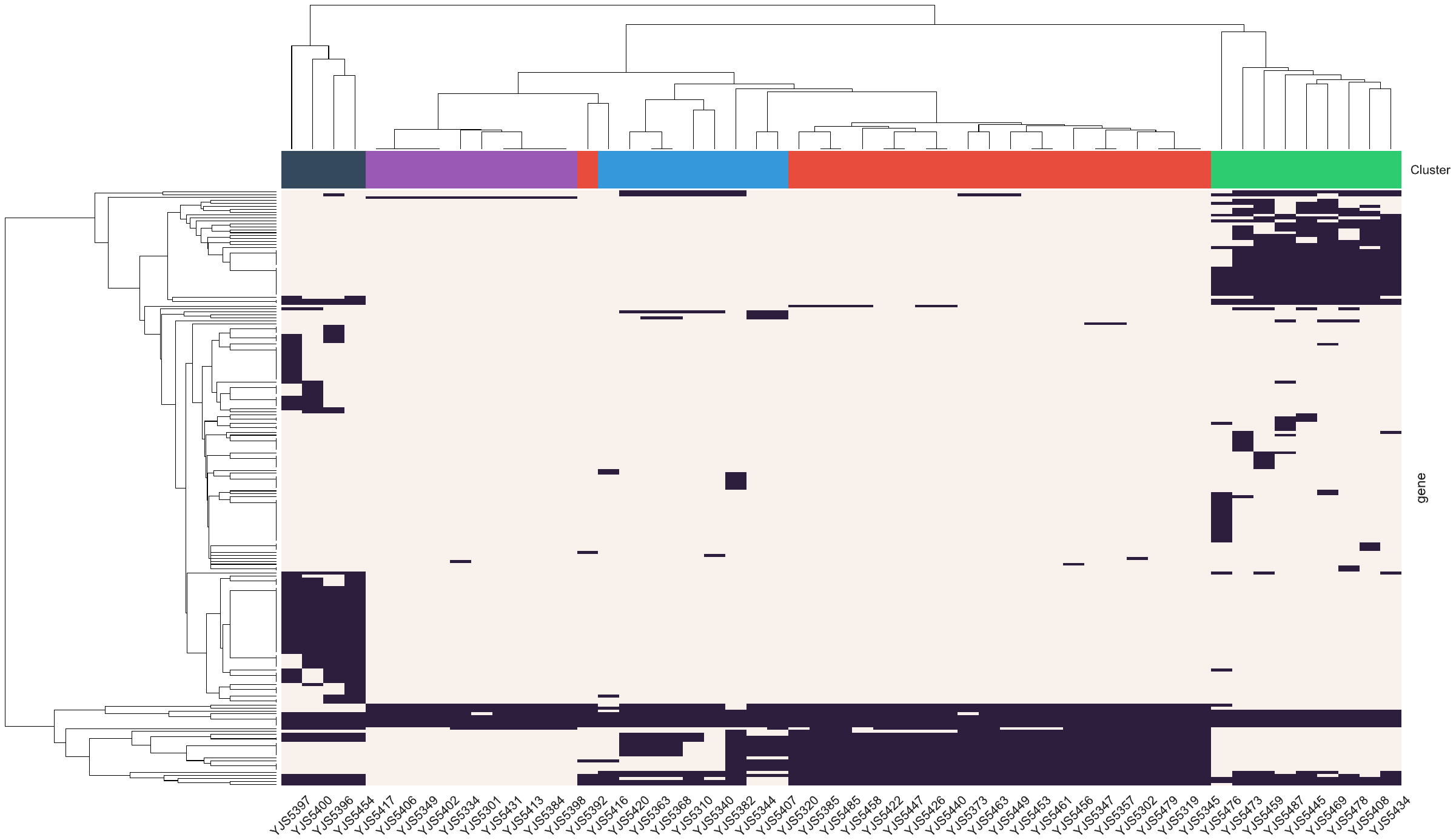

**Figure S8**. Heatmap of the 203 additional protein-coding genes within the population. Presence of the ORFs in a strain is indicated in dark purple. Both genes and strains were clustered using an euclidian metric with an average linkage. Colors on top represents the clusters as defined in Figure 1.
